# Expression Levels of the Attachment Protein G Differ Between Strains of a Murine Pneumovirus and Determine the Virulence

**DOI:** 10.64898/2026.01.22.701061

**Authors:** Akinbami R. Adenugba, Patrick Bohn, Jiangyan Yu, Markus Fehrholz, Anke K. Bergmann, Redmond P. Smyth, Christine D. Krempl

**Author notes:** Address correspondence to Christine D. Krempl.

## Abstract

Pneumonia virus of mice (PVM), the mouse homolog to respiratory syncytial virus (RSV), is increasingly used as surrogate model to study pneumovirus pathogenesis in a more natural pathogen-host relation. Two major strains of PVM, strain 15 and J3666 are currently used in laboratories, with preferences for either one or the other based on the well-documented isolation history of strain 15 or the suggested higher virulence of strain J3666.

Using conventional and long read sequencing, we found that the PVM strain J3666 represents two distinct virus populations, which are defined by sequence and structure of the G and SH genes encoding the putative attachment and small hydrophobic proteins, in addition to further nucleotide polymorphisms. Specifically, a nucleotide polymorphism at position 65 in the G gene results in either an upstream open reading frame (uORF) preceding the main ORF in frame, or an extension of the major G ORF by 18 codons. The impact of the different forms of the J3666-G genes on PVM was examined by generating recombinant PVMs differing exclusively in the distinctive 5’ portion of the respective G gene. This revealed that the population expressing a G protein with an extended main G ORF was more virulent, whereas the presence of a uORF attenuated virulence. The virulence of PVM correlated with increased expression levels of G, whereas attenuation was rather associated with downregulated expression of G due to the presence of a uORF. Thus, modulation of G protein levels may be an important mechanism by which pneumoviruses modulate virulence.

**Importance:** The pneumonia virus of mice strain J3666 is considered a more virulent and more suitable model for severe lower respiratory tract infections. The organization of the gene for the attachment protein G is reported to contain a small upstream open reading frame (uORF) preceding the main G ORF in frame. The translated G protein is predicted to comprise 396 amino acids. We report that this virus strain may be a mixture of two different populations, each with differing virulence. The more virulent population encodes a G protein of potentially 414 amino acids instead of a small uORF. This G gene organization is associated with an increased G protein expression. Importantly, this organization of the G gene is in line with that of several newly identified pneumoviruses, i.e., canine and swine pneumoviruses. These viruses may comprise a distinct group within the *Pneumoviridae* family.

## Introduction

Pneumonia virus of mice (PVM), with the official taxonomic name *murine orthopneumovirus*, is the first identified member of the genus *Orthopneumovirus* within the family *Pneumoviridae* (1, 2). An increasing number of laboratories are using the virus as a convenient experimental model to study acute respiratory disease caused by the respiratory syncytial virus (RSV) within a natural host (3). Two strains, 15 and J3666, have been characterized and are currently used in laboratories. A third independent strain, named strain Y, has been described in literature (3, 4). In addition, murine orthopneumovirus-like sequences were identified in bats or during an approach to characterize the virome of game animals in China (5, 6). Further orthopneumoviruses, i.e. canine, feline and porcine isolates (CnPV, FePV, SOPV), appear to be more closely related to the murine orthopneumoviruses than to RSV (7-9).

The PVM strain 15 is one of the original isolates described by Horsfall and Hahn (2). Its history is thoroughly documented in the scientific literature, thereby establishing it as reference strain. An avirulent variant, also designated as PVM 15/Warwick, resulting from continuous passage in tissue culture and plaque purification, exists in parallel to the original virulent isolate (3, 10, 11). In contrast, the isolation history of PVM strain J3666 is somewhat nebulous. The virus was first mentioned in 1995 by Easton and colleagues (10), and has subsequently been reported to originate from the same laboratory as strain 15, eventually even from the same isolate (3). The virus is reported to have a passage history predominantly in mice, although amplification of virus stocks also involved propagation in tissue culture (12, 13). As a result of its passage history, the latter one has been suggested to be more virulent and, for this reason, is preferentially used by several laboratories (3).

The sequences of the complete genomes of PVM strain 15, both the virulent and attenuated variants, and of strain J3666 have been described (12, 14). The single-stranded negative-sense RNA genomes contain 10 genes that potentially encode 12 proteins, in contrast to 11 proteins encoded by the RS viruses. In addition to the main open reading frame (ORF) the PVM-P mRNA encodes a second internal ORF that codes for a potential protein consisting of 137 amino acids. This protein is lacking a counterpart in the RS viruses, whereas the other 11 PVM-encoded proteins appear to be functional homologs to the RSV proteins (15-18).

The pneumoviruses encode three transmembrane surface glycoproteins, namely the small hydrophobic viroporin SH, the fusion protein F and the attachment protein G (19). The G proteins of pneumoviruses are type II transmembrane proteins, characterized by an N-terminus located within the virus particle. The ectodomain, which includes the C-terminus, comprises two-thirds of the polypeptide and is highly O-glycosylated, similarly to mucins. This overall structure, i.e. the percentage of potential O-glycosylation sites, is conserved. For RSV, the nucleotide and amino acid sequences are highly variable, such that e.g. subgroups and genotypes are defined by the sequence of G. In a similar vein, the most notable differences between PVM variants and strains, respectively, appear to reside in the organization of the G gene. The G gene of the reference strain 15 is a 1333-nucleotide sequence that encodes a G protein of 396 amino acids. This G protein is considerably larger than its RSV homologue, which has 282 to 319 amino acids, depending on the isolate (14). The main G ORF contains two potential in-frame start codons at positions 83 and 182, respectively, preceded by a non-translated region of 72 nucleotides. In the case of PVM strain J3666, the 5’ non-translated region appears to be divided by a small upstream open reading frame (uORF) starting with nucleotide 29 (AUG29) and coding for a potential peptide of 12 amino acids (10, 12). The start codon and uORF would be in-frame with the main G-ORF. A second form of the PVM J3666-G gene, which potentially coexists within the same virus stock, has been described. This form is marked by a nucleotide substitution that removes the stop codon of this uORF. Consequently, the substitution results in an extension of the main ORF by 54 nucleotides, which now encodes a potential G protein of 414 amino acids (11).

The function of the pneumovirus-G protein as an attachment protein is best characterized in the case of RSV. It is thought to mediate the initial binding to the target cells, i.e. ciliated cells, in the airway epithelium via interaction with the chemokine receptor CX3CR1. This primary interaction appears to be essential *in vivo* but not for infection of immortalized cell lines. It has been spontaneously deleted during propagation in cell lines or by directed genetic manipulation without major effect on replication in tissue culture with some dependence on the cell line (20-22). However, the replication of these mutants was found to be significantly attenuated in mice, and a vaccine candidate containing a G deletion proved to be overattenuated in children (20, 22). These findings point to an important role as virulence factor *in vivo*. These observations are replicated by corresponding PVM mutants, thus indicating that the PVM G protein has an *in-vivo*-function analogous to that of the RSV-G protein (18).

The present study revisits the potential coexistence of two G gene forms within the same PVM J3666 stock and identifies this strain as a mixture of two distinct populations. To analyze how the various G gene forms contribute to pathogenicity, the G gene of recombinant PVM 15 (rPVM) was replaced with either form described for PVM J3666. The presence of the uORF in the G gene reduced the pathogenicity of rPVM, which appeared to correlate with reduced G protein expression. In contrast, extending the major G ORF increased the expression of the G protein as well as the virulence of rPVM. Thus, the heightened virulence of strain J3666, as stated in the literature, may be attributed to the population expressing a G protein with a potentially extended cytoplasmic tail. Conversely, the presence of a uORF attenuates G expression and the virulence. Additionally, the differing pathogenicity of the viruses appears to be determined by the expression levels of the G protein.

## Results

### PVM J3666 is a mixture of two distinct G gene populations

We previously reported a potential nucleotide variation at position 65 of the PVM strain J3666-G gene that would either introduce a stop codon (nucleotide 65A, negative-sense RNA) so that the G gene would contain an in-frame small upstream open reading frame (uORF) preceding the main G ORF, as was originally described for PVM J3666, or code for lysine (nucleotide 65U, negative-sense RNA) leading to a 54-nucleotide extension (18 codons) of the main G ORF (10, 11). Considering the elapsed time and additional passages of the PVM stock, we decided first to reconfirm our previous observation in our present PVM J3666 stock.

To this end, a fragment covering part of the M gene, the complete SH and G genes, and part of the F gene was amplified from total cellular RNA of infected cells by RT-PCR using a proofreading polymerase. Direct sequencing of the G gene within the fragment confirmed our previous findings regarding nucleotide variations at position 65 (A or U in negative-sense RNA) (Fig. 1A). Moreover, all other previously observed nucleotide substitutions were confirmed: nt 104 (4614 in the genome) C for U (Gly for Ser), 165 (4675 in the genome) A for C (Val for Gly) and 1121 (5631 in the genome) U for A (Thr for Ser). However, we noted underlying peaks similar to those observed for position 65, indicating nucleotide variations also for these positions (data not shown).

**Figure 1.**
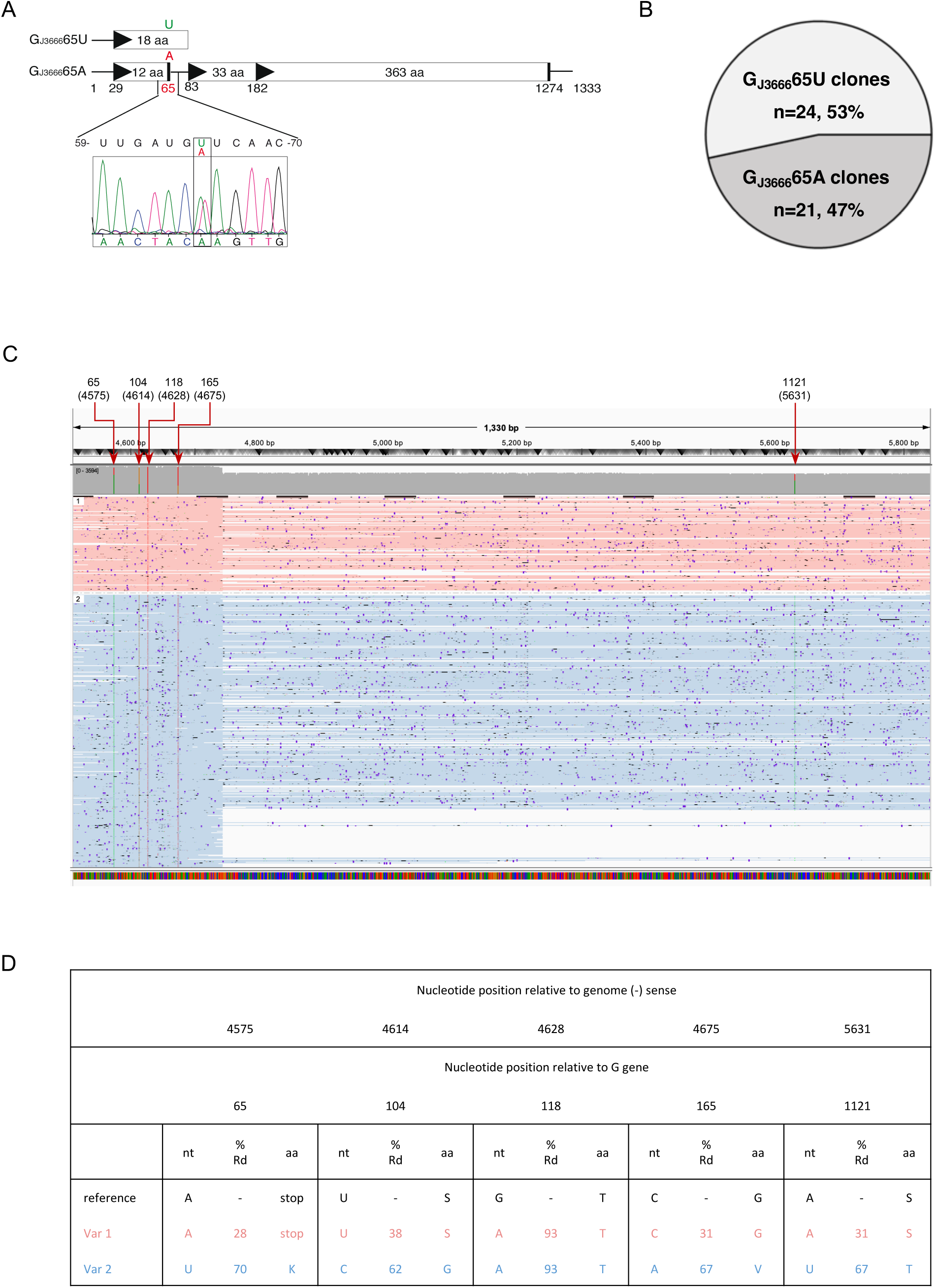
Analysis of the nucleotide 65 polymorphism of the PVM strain J3666-G gene. Total cellular RNA from PVM J3666-infected BHK-21 cell was subjected to RT-PCR amplifying a PVM fragment of 2,517 bp that covered 406 nt of the M gene, the entire SH and G genes and 314 nt in the F gene. The consensus sequence was determined by direct sequencing of the amplification product. (A) Schematic representation of the G gene and the nucleotide polymorphism at position 65 with consequences on ORFs. ORFs are depicted as open rectangles with potential start codons indicated by filled triangles and stop codons by a bar. The first and last nucleotides of the gene, and first nucleotides of the translational start and stop codons are indicated. Below the drawing, a representative electropherogram of the sequencing reaction for position 59 to 70 of the G gene is shown. Note that sequence data are in positive sense, whereas the genomic (-) sense RNA sequence is shown above. The overlaying peaks for nt 65 representing the first nt of the codon (underlined) for translational stop (UAG) and lysine (AAG), respectively, are boxed. (B) Percentage distribution of the sequence variation at position 65 within the PVM J3666 preparation. To analyze the percentage composition of the population, the RT-PCR fragments were cloned and the G gene sequences of 45 individual clones were determined. (C) Confirmation of sequence variation by ONT sequencing. The haplotagged reads spanning the full G gene (position 4511-5843 of the reference genome AY753909) are visualized using the Integrative Genomics Viewer (IGV). Reads corresponding to Var1 and Var2 are shown in pink and blue, respectively. Nucleotide polymorphisms relative to the gene and genome (in brackets) are pointed out by arrows. (D) Percentage distribution of nucleotide variations and amino acid exchanges according to read-based phasing.

To determine the percentage distribution of these nucleotide variations, the described RT-PCR fragments were inserted into plasmids and sequenced. Overall, six distinct cDNA preparations were generated from three independent RNA purifications, and the cloned PCR fragments were obtained from these six preparations. Of the 45 clones analyzed, 24 clones (53%) contained a G gene with a single extended G ORF starting at AUG29 (nt 65 U, negative-sense RNA), whereas 21 clones (47%) contained a G gene encoding a small uORF (nt 65 A, negative-sense) preceding the main G ORF that starts at AUG83 (Fig. 1B). Analysis of the clones with respect to other variable nucleotide positions revealed that clones encoding an A at G gene position 65 would encode a U at position 104 (Table S1). In addition, 17 of these clones were analyzed up to position 165, where C was encoded in all cases. In comparison, the clones that encoded U in position 65 with a single exception all encoded C in position 104, and 20 out of 23 analyzed up to position 165 encoded U in this position.

Long-read sequencing methods, such as Oxford Nanopore sequencing (Oxford Nanopore Technology, ONT), enable direct characterization of virus populations by sequencing entire viral genomes in single reads. To confirm the results obtained by conventional Sanger sequencing, we applied ONT sequencing to the virus stock used for previous experiments. We designed a sequencing strategy that enabled us to phase single nucleotide polymorphisms across the whole PVM genome. Specifically, we designed three PCRs, two of which overlapped the SH-G region. We selected the highly processive reverse transcriptase MarathonRT and Primestar GXL polymerase for cDNA synthesis and amplification, and performed long read DNA sequencing with Kit 14 on a Flongle and MinIon R10.4.1 flow cell.

Overall, 16,024 reads (49 megabases) were generated, which matched the expected amplicon sizes (Fig. S1) and could be aligned to the reference sequence NC_006579 of PVM strain J3666. The mean sequencing depth of the G gene was 2,831x. The data confirmed the polymorphism at position 65 of the G gene (i.e., 4575 of the genome), with 28% of reads indicating A and 70% of reads indicating U (Fig. 1C-D). Furthermore, differences in position 104 (4614 of genome), 165 (4675 of genome) and 1121 (5631 of genome) when coding for U in position 65 were confirmed (Fig. 1C-D). This almost homogeneous distribution of nucleotide polymorphisms pointed to two discrete populations that we preliminarily termed G_J3666_65A and G_J3666_65U based on the originally described variation.

Analysis of the full genomic sequence revealed further nucleotide polymorphism compared to the published sequence in seven out of ten genes (Fig. 2A). The 5’ non-coding region of the M gene, the SH gene, and the first 200 nts of the G gene presented itself as a hotspot of nucleotide polymorphism, as summarized in Fig. 2B and Supplemental table S2. Overall, 34 nucleotide polymorphisms distributed throughout the M and SH genes, in addition to the ones already described for G, were identified. Most of these nucleotide polymorphisms were specific to either the G_J3666_65U or G_J3666_65A genomes, thus confirming the presence of two defined populations within PVM strain J3666 (Fig. 2B). Of note, 27 of the nucleotide exchanges involve an A to G (- sense) replacement from the reference sequence to the J3666-65A population, which is the most related sequence, but not to J3666-65U. This could point to a single hypermutation event as mediated by the adenosine deaminase acting on RNA (ADAR) 1 on vRNA (A to I, i.e. U to C on cRNA). Five of these putative hypermutations are in the M gene end signal and the SH gene start signal, respectively, which potentially affects their functionality (Table S2). Furthermore, 16 nucleotide exchanges in the SH ORF also cause animo acid substitutions.

**Figure 2.**
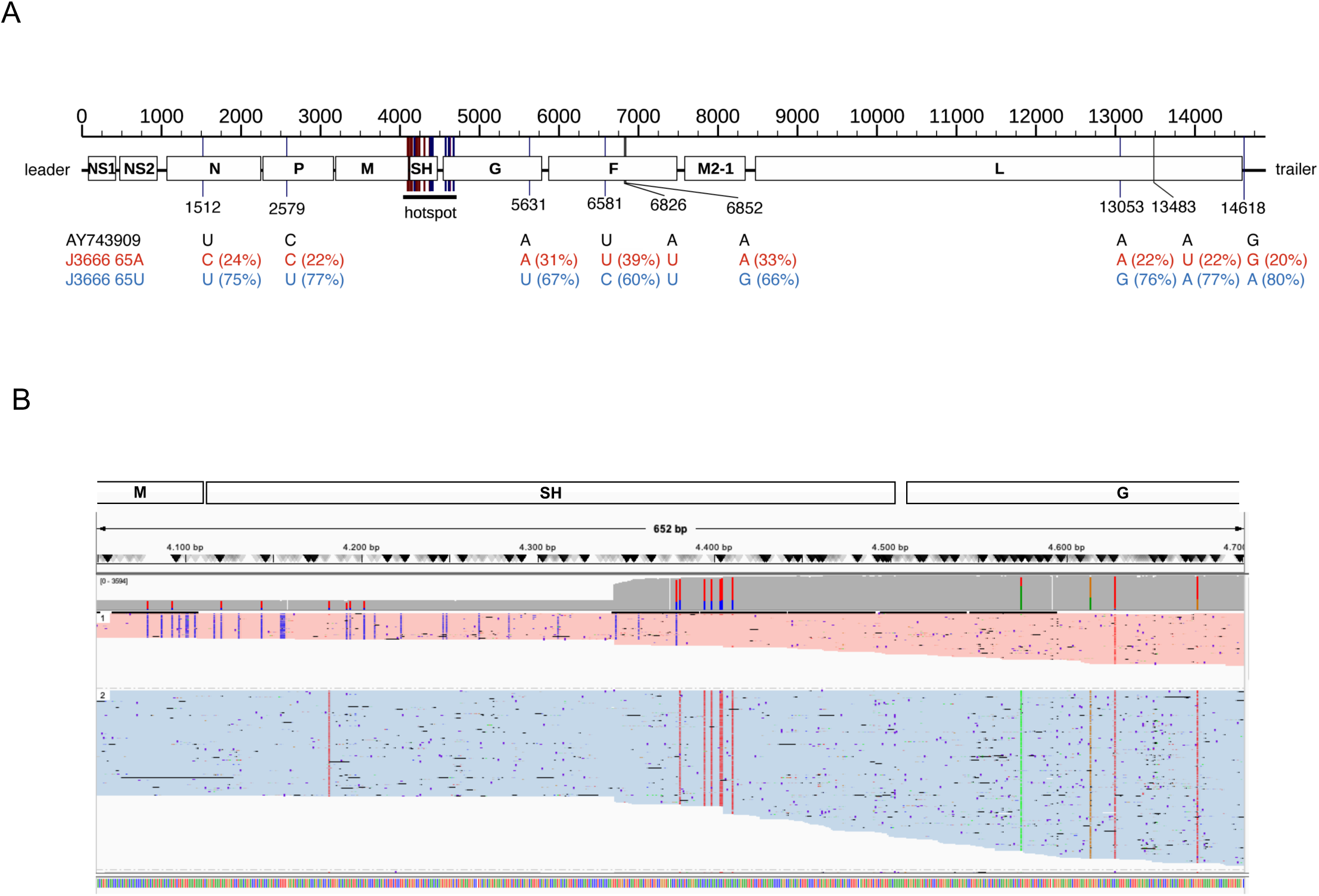
Nucleotide polymorphisms distributed over the full genome divide PVM J3666 into two discrete populations. (A) Schematic representation of the PVM J3666 genome with positions of polymorphisms, including the hotspot covering the 5‘ noncoding part of the M gene, the entire SH gene, and the first 200 nts of the G gene. Except for the hotspot, the nucleotides in the reference sequence, as well as the polymorphisms aligning with the genomic 65A or 65U nts, are indicated below the scheme along with the percentage distribution of reads. (B) IGV visualization showing the accumulation of nucleotide polymorphisms in the region from nt 4052 to 4700 of the PVM J3666 genome, in the two discrete populations. The colors of the reads correspond to those presented in Figure 1C-D. The respective genes are indicated above the ruler.

Of the seven remaining nucleotide substitutions that represent G to A (-sense) changes from the reference sequence to the J3666-65U population, the most prominent effect would result in the expression of differentially sized SH proteins, with the A (- sense) at position 287 of the SH gene (nt 4399 of the genome) representing the first nucleotide of the stop codon, thus ending the SH ORF after 92 codons, as opposed to G at this position, which would induce a codon for glutamine.

Taking our results together, we conclude that PVM J3666 represents a mixture of two distinct virus populations, which are mainly represented by the sequence, organization and ORF sizes of the G and SH genes.

### Recombinant PVM differing exclusively in the organization of the G genes

Since major differences in the pathogenicity of PVM were previously attributed to the G protein (11, 18), we focused on the variability of the PVM J3666-G gene. To investigate the functional differences between the two gene variants, recombinant PVM (rPVM) based on the PVM 15 backbone were generated that would encode the G gene containing a uORF (rPVM-G_J3666_65A) or encode a potentially extended G protein (rPVM-G_J3666_65U), respectively. For generation of rPVM-G_J3666_65U, an *Age*I site that is unique for rPVM was added upstream of the G_J3666_65U gene by PCR amplification of an appropriate G clone described above. Subsequently, a segment of the strain 15-G gene, encompassing nt 1 to 925, was substituted with the previously amplified fragment of G_J3666_65U (Fig. 3A). Thus, the resulting chimeric G gene also contained the nucleotide substitutions at positions 104, 165 and 1121, which are specific for the G_J3666_65U population, since the latter nucleotide is identical with that of strain 15. The generation of rPVM-G_J3666_65A involved the addition of an *Age*I site to G_J3666_65A, similar to the previously described method. In this instance, a segment of PVM 15 containing the complete G gene and 155 nucleotides of the adjacent F gene was replaced with a PCR fragment of the J3666-65A subpopulation. Notably, there are no sequence differences in the G-F intergenic region or the first 155 nucleotides of the F gene between strains 15 and J3666. Thus, despite the use of diverse cloning strategies, the resulting rPVM mutants differed exclusively in their G genes, with the complete respective G gene equaling that of the J3666 subpopulations. The parental rPVM strain 15 has been described previously and was used for comparison (18).

**Figure 3.**
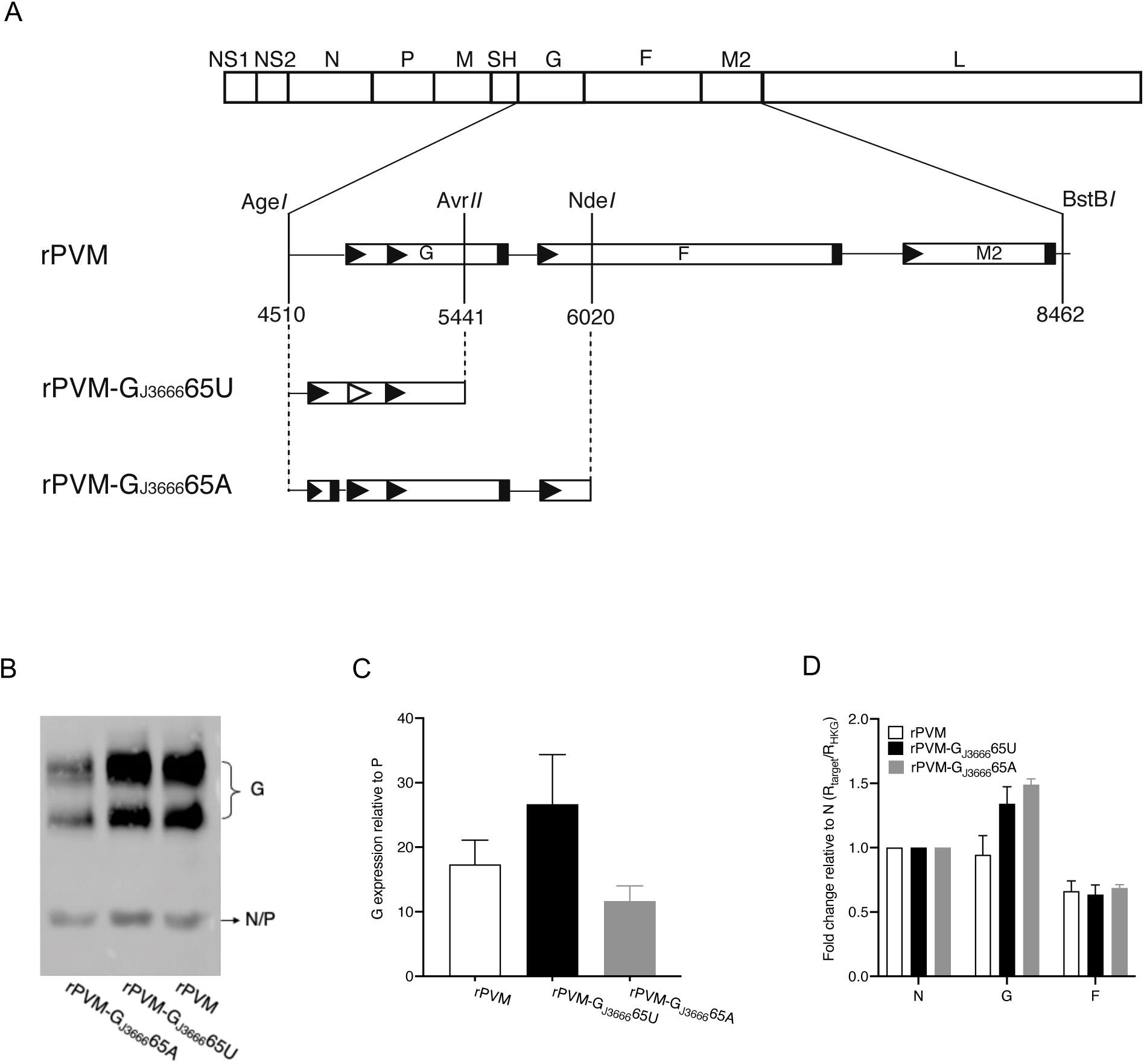
Recombinant PVM with G genes corresponding to J3666 65A or 65U, respectively. (A) Schematic representation of rPVM encoding the two G gene variants of strain J3666. All changes were performed using a plasmid containing an antigenomic cDNA fragment encoding the G, F, and M2 genes, which was flanked by unique *Age*I and *BstB*I restriction enzyme sites for reassembly of the antigenomic full-length plasmid. Fragments of J3666-G containing the 65U or 65A nts, as well as other variant-determining nts, were amplified from plasmids using a forward primer that added an *Age*I restriction site specific for rPVM. The original G strain 15 fragments, as indicated in the drawing, were exchanged for corresponding G_J3666_65U and G_J3666_65A fragments. Nucleotide sequences between positions 4510 and 6020 of the genomic sequence were confirmed by sequencing. Nucleotide positions of restriction enzyme sites are given relative to the rPVM genome. (B - D) G protein expression by the rPVM-G gene variants. (B) The expression of G protein was determined in cell lysates collected from BHK-21 cells infected with rPVM, rPVM-G_J3666_65A, or rPVM-G_J3666_65U, respectively, by western blotting. PVM proteins were detected by immunostaining using serum from a PVM-infected convalescent mouse, combined with chemiluminescence imaging using the LAS 3000 CCD camera (Fujifilm). (C) Chemiluminescence signals from two independent experiments were quantified using the AIDA software (v.3.20.116, Raytest, Berlin). For each lane, the amount of G isoforms per lane were expressed as the ratio of P. The standard error between runs is represented by error bars. (D) The transcription rate of the G, F and N genes was determined by SYBR green-based qRT-PCR. Total cellular RNA was isolated from BHK-21 cells infected as described above, reverse transcribed with random hexamers, and subjected to qPCR with primers specific for the indicated mRNAs. The data were analyzed with the LinRegRCR program (37). The result for each target gene was normalized to 18s rRNA and expressed as fold change relative to the amount of N mRNA. Error bars represent the standard deviation between two independent runs each performed in triplicate.

To characterize the recovered rPVM-G variants, we examined the expression of the G proteins. BHK-21 cells were infected with rPVM-G_J3666_65A, rPVM-G_J3666_65U, or parental rPVM, and G protein expression in cell lysates was analyzed by western blotting. The amount of G protein was quantified relative to the expression of P protein by chemiluminescence imaging. To exclude any transcriptional effects on G protein expression, mRNA levels of G, an upstream gene (N), and a downstream gene (F) were determined by quantitative RT-PCR. The results were normalized to the amount of 18S rRNA and analyzed relative to the intrinsic N mRNA levels for each virus.

This analysis revealed that the presence of the uORF moderately reduced the expression of the G protein from the G_J3666_65A gene by 1.8-fold (or 20%), compared to the relative amount of G protein expressed in cells infected with rPVM (Fig. 3B-C). Unexpectedly, the amount of G protein in lysates of BHK-21 cells infected with the rPVM-G_J3666_65U variant increased on average 1.5-fold when compared to the amount of G protein in rPVM-infected cells. These differences in protein expression were not due to differing transcription levels, as the ratio of G/N mRNA ratio was comparable, ranging from 0.9- to 1.5-fold, although it was slightly more variable than the F/N mRNA ratio (Fig. 3D).

An increase in the size of the G_J3666_65U gene-derived protein was not observed, although 18 amino acids corresponding to 2.1 kD may have been acquired due to the extension of the main ORF. This is not unexpected since a truncation of the cytoplasmic tail for 33 amino acids could not be detected by electrophoretic mobility in SDS-Page (18). It is likely that the size differences are obscured by the extensive glycosylation of the G protein. In summary, deleting the uORF stop codon, which extends the main ORF to AUG29 in G_J3666_65U, appears to enhance the expression of the G protein.

### The presence of a uORF in G promotes replication in BHK-21 cells

To determine the effect of the different G gene and ORF variants on replication in susceptible cell lines, multi-cycle replication kinetics were carried out using BHK-21 and macrophage-like RAW 264.7 cells infected with rPVM-G_J3666_65A, rPVM-G_J3666_65U, and rPVM, respectively, at an MOI of 0.01 PFU/cell. The virus yield was determined for days 1, 3, 5, and 7, respectively.

In BHK-21 cells, PVM expressing a potentially N-terminal extended G protein, rPVM-G_J3666_65U, replicated almost indistinguishably from rPVM, with each virus reaching titers of approximately 7 x 10^5^ PFU/ml by day 7 (Fig. 4A). Compared to the other two viruses, rPVM-G_J3666_65A replicated more efficiently, with titers 3-fold higher by day 3 and 6-fold higher by day 7. Of note, the presence of the uORF in G_J3666_65A represents the major difference from the G gene of the parental rPVM.

**Figure 4.**
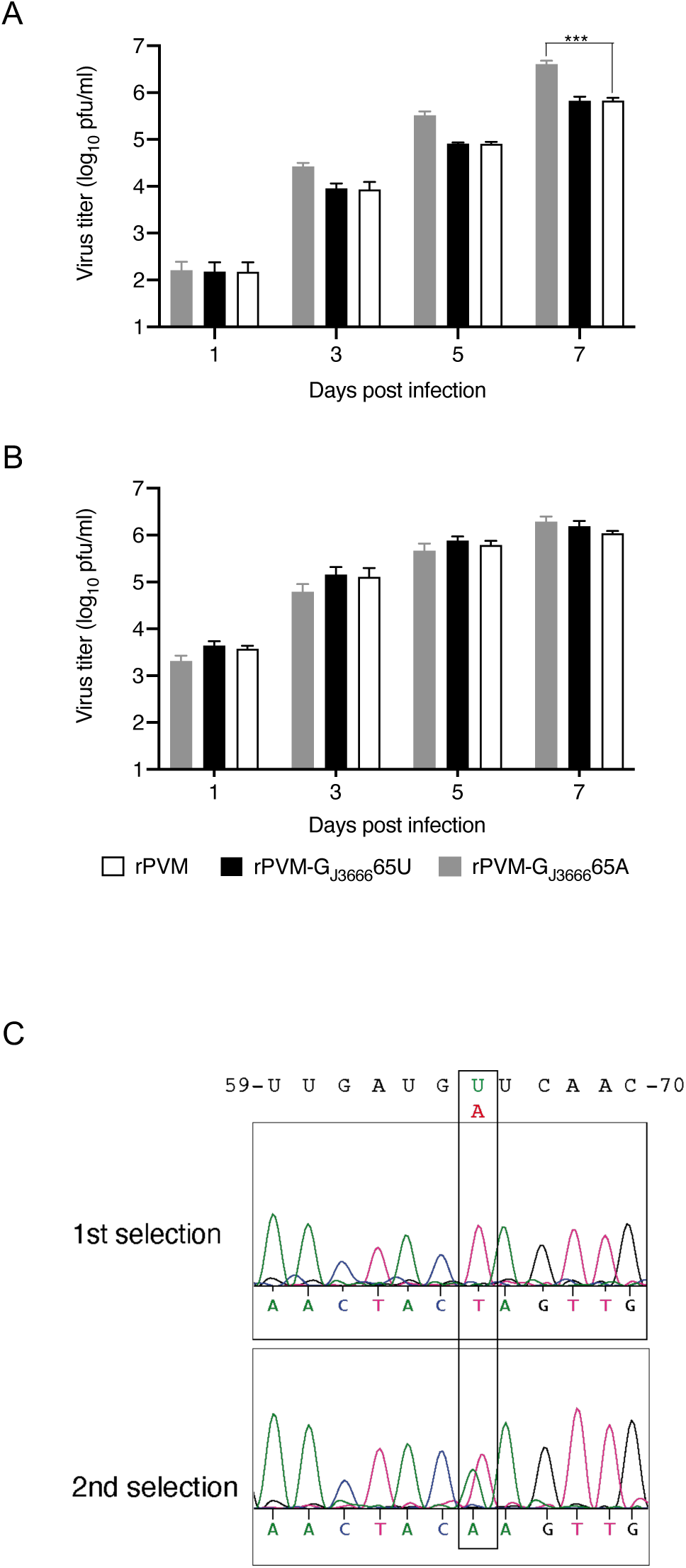
Replication kinetics of PVM-G gene variants. (A) BHK-21 or (B) RAW 264.7 cells were infected with rPVM, rPVM-G_J3666_65U, or rPVM-G_J3666_65A at an MOI of 0.01 PFU/cell. At the indicated time points, the cells and their supernatants were harvested, subjected to freeze-thaw cycles to detach cell-associated virus particles, clarified from debris, and stored for determination of virus titer by plaque assay. Data points represent mean values of two independent experiments performed in duplicate. Standard error is represented by error bars. Statistical analysis was carried out using 2-way ANOVA: for the Bonferroni post-hoc test, rPVM-G_J3666_65A and rPVM-G_J3666_65U were compared to rPVM as the control. Differences were considered significant at p < 0.05 (***, p ≤ 0.001). (C) Sequence analysis of the PVM J3666 stock following two individual *in vitro* selection experiments by sequential passaging in BHK-21 cells. Electropherograms of the sequencing reactions for nts 59 to 70 of the G gene after seven (1^st^ experiment) and eight passages (2^nd^ experiment), respectively, are shown. Sequence data are in positive sense, whereas the genomic negative-sense RNA sequence is shown above. Peaks for nt 65 are boxed.

To obtain data from a second cell line susceptible to PVM, the replication kinetics were also determined using RAW 264.7 cells. In this murine macrophage-like cell line, all three PVM-G variants replicated with comparable kinetics, reaching similar titers at all time points investigated (Fig. 4B).

In order to evaluate if propagation in cell cultures may favor one of the two populations, particularly one or the other G gene variation, the mixed population was serially passaged in BHK-21 cells in two separate approaches. Following each passage, the SH and G genes were amplified from total cellular RNA of infected cells, and the region covering nucleotide 65 of the G gene was sequenced. In one experimental approach, the sequencing revealed a homogeneous G_J3666_65A population following seven passages that was confirmed by analysis of single clones, thus, indicating a slight advantage of a uORF for replication in BHK-21 cells; however, in a 2^nd^ experiment, eight serial passages did not select for one of the two G gene populations (Fig. 4C).

Thus, the presence of a small uORF seems to favor replication in some cell lines, which is consistent with the selection for this variant upon sequential passages in BHK-21 cells, while the extension of the G ORF has no effect on replication efficiency.

### The organization of the G gene affects the pathogenicity in mice

Lastly, to examine the effect of the varying G_J3666_-gene organization on replication and the virulence of PVM *in vivo*, BALB/c mice were infected intranasally with 150 PFU per mouse of rPVM-G_J3666_65A, rPVM-G_J3666_65U, or the parental rPVM. Disease was evaluated using weight loss as an objective assessment method. To determine replication efficiency, groups of mice were sacrificed on days 3 and 6 post-infection, respectively, the lungs were removed, and the virus load in lung homogenates was determined by plaque assay.

As expected, infection of mice with rPVM resulted in mild transient weight loss by a maximum of 6% over two days, starting after day 7, followed by full recovery until day 12 (Fig. 5A). This weight loss was accompanied by apparent signs of disease, i.e., ruffled fur and reduced activity (data not shown). On day 3, the average pulmonary virus load was 5.7 x 10^2^ PFU/lung, and on day 6, it was 3.8 x 10^5^ PFU/lung (Fig. 5B). In comparison, mice infected with the equivalent dose of rPVM-G_J3666_65A did not lose weight or show any clinical symptoms throughout the duration of the experiment, indicating that the presence of a uORF somewhat attenuated the virulence of PVM. This attenuation appeared to be independent of replication efficiency, since pulmonary titers of rPVM-G_J3666_65A were not significantly different from those of the parental virus on both days determined (7.5 x 10^2^ PFU/lung vs. 5.7 x 10^2^ PFU/lung on day 3; 3.7 x 10^5^ PFU/lung vs. 3.8 x 10^5^ on day 6). In marked contrast, the weight loss of mice infected with rPVM-G_J3666_65U was more pronounced, exceeding 10% at the peak of disease (Fig. 5A). The enhanced virulence of rPVM-G_J3666_65U was accompanied by a virus load that was moderately increased, i.e. 2-fold, compared to that of rPVM-infected mice at day 3. However, the virus load leveled out at 5.5 x 10^5^ PFU/lung for rPVM-G_J3666_65A and rPVM at day 6, the peak of virus replication. Thus, the G_J3666_65U variation appeared to provide a slight replicative advantage *in vivo* at early time points of infection.

**Figure 5.**
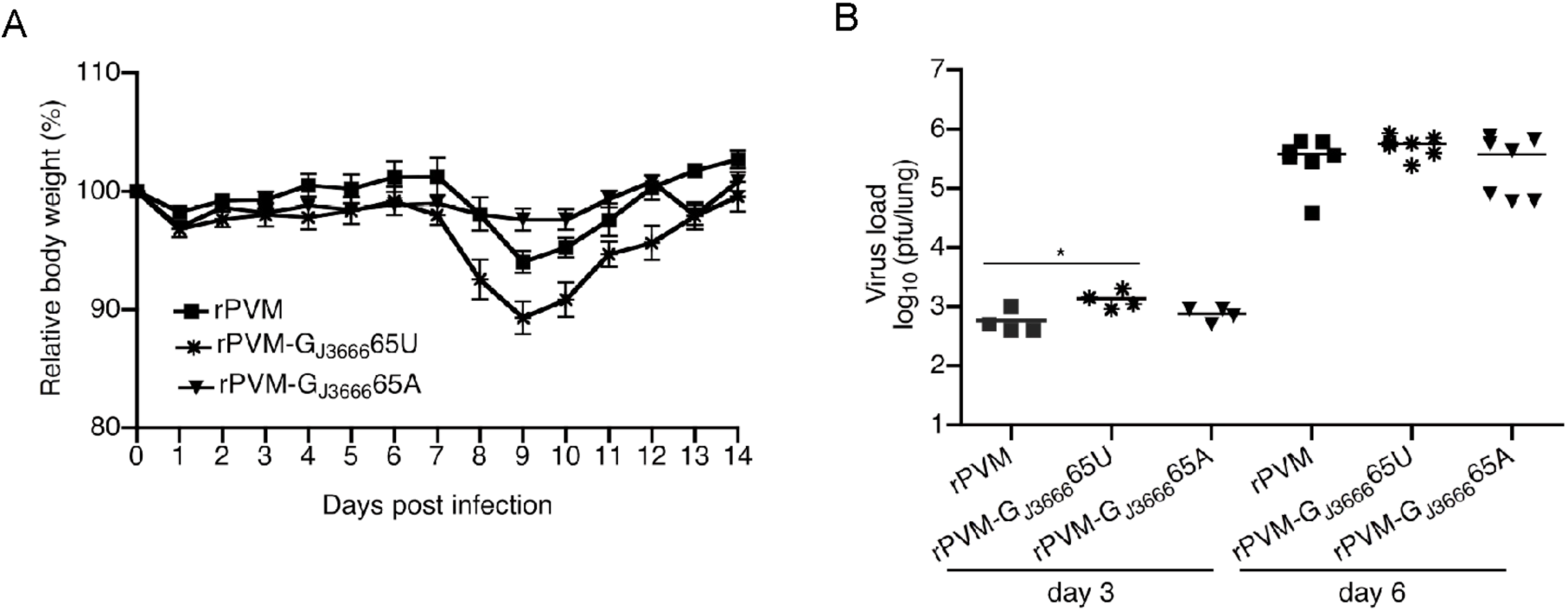
Virulence and replication of rPVM-G gene variants in the respiratory tract of BALB/c mice. Mice in groups of 5 were infected intranasally with 150 PFU of rPVM, rPVM-G_J3666_65A or rPVM-G_J3666_65U, respectively, in a volume of 80 µl. (A) For comparison of virulence, the mice were observed closely and the body weight determined on a daily basis. Results represent mean values with standard error of the mean derived from five mice (rPVM-G_J3666_65A or rPVM-G_J3666_65U), respective 4 mice (rPVM) per group. (B) For determination of replication, mice were infected as described for panel A and sacrificed at the indicated days after infection. The lungs were removed, homogenized and the virus load was determined by plaque assay. Statistical analysis was performed using the Krusal-Wallis test, and rPVM-G_J3666_65A and rPVM-G_J3666_65U were compared to rPVM. The differences were considered statistically significant at P ≤ 0.05 (*, P ≤ 0.01).

In summary, the two G gene variants reported for PVM J3666 have the capability to confer a differing pathogenicity phenotype to the respective virus subpopulation. Interestingly, the level of pathogenicity correlated with the level of G protein expression, i.e. G_J3666_65U over G strain 15 over G_J3666_65A, thus indicating an association *in vivo*.

## Discussion

Two PVM strains, 15 and J3666, are currently used as experimental model to study the pathogenesis of pneumoviruses in a natural host. Preferences of investigators for either one or the other may be based on the well documented isolation history of strain 15, or the suggested higher virulence of strain J3666 due to frequent mouse passages (3, 10, 13). In the present study we found that what was thought to be PVM strain J3666 actually represents a mixture of two distinct virus populations mainly distinguished by the sequence and structure of their G and SH genes. Moreover, focusing on the G protein as a major virulence factor we found that the two populations differ in the expression levels of the G protein that confer a differing virulence.

The existence of two distinct populations was first analyzed by conventional methods that involve subcloning and sequencing of RT-PCR fragments. According to the reference sequence ((12), NC_006579.1), the J3666-G gene would encode two open reading frames, i.e. the main G ORF starting at AUG83 and coding for 396 amino acids that is preceded by a small uORF starting at AUG29 and potentially coding for a peptide of 12 amino acids. In the subpopulation described in the present study the main G ORF starts with AUG29 and codes for a polypeptide of 414 amino acids. The co-existence of two distinct PVM populations was confirmed by ONT sequencing this time covering the full genome (Fig. 1C, Fig. 2) that revealed a roughly 30% to 70% distribution in favor of the population with the G ORF starting at AUG29. According to this, the population expressing a G protein of potentially 414 aa represents the major population which corresponds to our previous data (11).

Additional population defining features are found in the SH gene, the most characterizing one introducing a preterminal stop codon in the population coding for a larger G protein. This preterminal stop codon causes shortening of the SH protein for 22 codons from 114 aa to 92 aa compared to the reference sequence. We observed overall 34 polymorphisms in the M and SH genes, five of them involving the M-gene end and the SH-gene start signals, respectively, identifying this region as a hot spot. Most of these polymorphisms were not reported in the reference sequence, are specific for the subpopulation expressing a G of 396 aa and appear be hypermutations that may result from a single event. When and where this event occurred is not clear. However, eight of the 27 polymorphic positions of the SH gene identified by us, i. e. nucleotides 44, 70, 269, 283, 287, 292, 293, and 299, had also been reported to be variable independently from us and was accounted for the quasispecies nature of viruses (12). This also included a size plasticity of the SH protein for 92, 96 or 114 amino acids. Thus, this heterogeneity of strain J3666 is not likely to be specific for the virus preparation used by our lab, whereas the percentage distribution may be.

In recent years further pneumoviruses of dogs, cats and swine have been identified that are more closely related to PVM than to the prototype orthopneumovirus human RSV (7-9, 23-31). Of note, the G genes of canine pneumoviruses (CnPV) and swine orthopneumoviruses (SOPV) are predicted to encode G proteins of 414 aa (Fig. S2) and an SH ORFs of 92 aa (Fig. S3), respectively, comparable to the J3666-population identified by us. In addition, the PVM Y strain, which was isolated independently, appears to encode a G of potentially 414 aa (Fig. S2). This differs from what was previously annotated (GeneBank ac No JQ899033). Thus, we suggest that a G gene that codes for a G protein of potentially 414 aa represent the defining G gene organization for this group of pneumoviruses.

PVM J3666 has been reported to be more virulent in mice than the reference strain 15 (ATCC) (32). However, this increased virulence is attributed to the published reference sequence (NC_0065799) that would contain a uORF preceding the main ORF in the G gene. By constructing recombinant PVM containing the different G gene forms we confirmed a heightened virulence of J3666 compared to strain 15. However, the more virulent form correlated with the G ORF starting at AUG29 and coding for a potential G protein of 414 aa, whereas the population containing a uORF preceding the main G ORF and coding for a G protein of 396 aa was the least virulent (Fig. 5). The virulence of strain 15, that encodes a G protein of 396 aa located in between. Of note, nucleotide 30G in the 5’-UTR of the G-strain 15 gene (- sense) ablates a canonical start codon that otherwise would be corresponding to AUG29 and be the first AUG of a major G ORF.

Interestingly, the pathogenicity of the different PVM-G variants correlated with the expression levels of the G protein in infected cells. The low-pathogenic PVM with the reference sequence G gene was characterized by reduced G protein expression compared to the parental recombinant strain 15 virus, likely due to the ribosomal diverting activity of uORFs (33). Conversely, the variant with a G ORF starting at AUG29 expressed the highest levels of G protein (Fig. 3 B, C). Notably, the predicted G proteins of strains 15 and J3666, which both start at AUG83, differ by only four amino acids, i.e L5F, V28G, T112A and T347S. When the main G ORF starts at AUG29, it is in a suboptimal context for translational initiation (CAA<u>AUG</u>A, AUG29 underlined) and is predicted to promote ribosomal leaky scanning, compared to the improved contextual strength of AUG83 (AUC<u>AUG</u>G, AUG82 underlined) (34). Consequently, the majority of translation would be predicted to start with AUG83, as does the G ORF of strain 15, and polypeptides initiated at AUG29 would add up to the majority of polypeptides initiated at the downstream AUG. Thus, it is more likely that increasing of G protein levels rather than extending of the cytoplasmic tail is causing the heightened virulence of the virus.

Intriguingly, the G gene of RSV also contains a uORF that overlaps the start codon of the main ORF. This uORF has been shown to downregulate the G protein’s overall expression, but without significance for replication in infected cell lines (22). Remarkably, this uORF is conserved in all hRSV. Thus, this uORF may play a role in controlling G levels and limiting the pathogenicity of RSV.

In summary, we propose that the virulent form of PVM J3666 strains encodes a G protein with an AUG29 start codon in the main ORF, as well as a 92 aa SH protein. These features appear to be typical of the most closely related pneumoviruses, such as CnPV and SOPV. Thus, we propose that these G and SH gene forms represent the natural form of PVM.

## Material and Methods

### Cells and viruses

The origin of PVM J3666 has been described (11) and had been passaged 8 times in BHK-21 cells since receipt. Propagation of PVM J3666, PVM strain 15 and recombinant pneumonia virus of mice (rPVM) as well as determination of virus titers by plaque assay has been described previously (18). Vero cells were cultivated in Eagle’s MEM supplemented with 10% fetal calf serum. BHK-21 cells and BSR-T7/5 cells constitutively expressing the T7 polymerase (35) were maintained in Glasgow’s MEM. RAW 264.7 cells were grown in Dulbecco’s MEM supplemented with 1X sodium pyruvate (Invitrogen) and 10% fetal calf serum.

Growth curves were performed in duplicate. BHK-21 cells RAW 264.7 cells seeded in 12-well plates were infected at MOI of 0.01 PFU per cell. Cells and supernatant were then collected at the indicated time points, cell-bound virus detached by vigorous mixing, cell debris removed by low speed centrifugation and aliquots were flash frozen and stored at −80°C until used.

Selection experiments were performed by sequential passaging of biological PVM J3666. For each passage, BHK-21 cells were infected at an MOI of 0.01 for 7 days, then cells and supernatants were harvested as described above. Cell debris was used for RNA isolation and supernatant was passaged on.

### Amplification and cloning of PVM fragments for sequence analysis

Total cellular RNA was collected from infected cells as described previously (14). The RNA was reverse transcribed with RevertAid H minus reverse transcriptase (Thermo-Scientific, Leon-Rot, Germany) and random hexamer primer (Qiagen, Hilden, Germany) followed by RNaseH treatment (NEB, Frankfurt, Germany). Thereafter, 5 µl of RT reaction were subjected to 35 cycles of PCR amplification using PVM J3666-specific primers (PVM M fw: 5’-<u>GGTACC</u>TTGATAGTGTGGAGCAG; PVM F rs: 5’ <u>GGTACC</u>AAGAATCAAACCGAGGAAC) and Phusion high fidelity polymerase (NEB, Ipswhich). Note that the primer added artificial *Acc65*I sites (underlined) to both ends of the amplificates for cloning into pGEM3zf(+) or pUC19 plasmids. Nucleotide sequencing was carried out using an ABI 3100 sequencer and the Big-Dye terminator mix version 1.1 (Applied Biosystems, Darmstadt,Germany).

### Quantification of PVM transcripts

Total cellular RNA was collected from BHK-21 cells infected at an MOI of 1 PFU/cell after 24 h with the RNeasy RNA Isolation kit (Qiagen, Hilden, Germany) as specified by the manufacturer, and 1 µg of the RNA was reverse transcribed as described above. The synthesized cDNAs were diluted 1:10 with H_2_O and quantified in triplicates using the iTaq universal SYBR Green supermix using following primer pairs: GACCGACCTGATTTACCT (PVM G fw), CACCATTGTTTAAGCCCA (PVM G rs); the primer pairs for N and F, and 18S rRNA are published (15, 36).

The specificity and sizes of the PCR products were confirmed with the melt curve and gel electrophoreses, respectively. The data were analyzed using the LingRegPCR software (37). The N, F and G mRNA were first normalized to their respective 18s rRNA, then the normalized F and G mRNA were analyzed to the normalized N mRNA.

### Nanopore sequencing of virus stock

Viral RNA of 160 µl of virus stock was extracted using the Macherey-Nagel Viral RNA mini kit. RNA was eluted in 20 ul elution buffer, followed by reverse transcription of 5 ul eluate in a 20 µl reaction containing 40 U MarathonRT in x1 RT buffer (50mM Tris-HCl pH 8.3, 200mM KCl, 20% (v/v) glycerol) and primers RT1, PCR A re and PCR B re. Marathon RT purified according to Zhao et al. (38); pET-6xHis-SUMO-

MarathonRT was a gift from Anna Pyle (Addgene plasmid # 109029; http://n2t.net/addgene:109029; RRID:Addgene_109029). After incubation at 42 °C for 2 h the RT was inactivated by heating to 80 °C for 5 min, followed by cDNA purification with the Macherey-Nagel PCR Clean-up kit using NTC buffer and elution in 12 µl elution buffer. Next, PCRs to generate amplicons A, B, C, BC and ABC were performed with Primestar GXL. Amplification success was confirmed on a 1% agarose gel in TAE buffer and amplicons were purified with 1 volume of SPRI beads (Omega Biotek MagBind NGS), washed twice with 80 % EtOH and eluted in 10 µl H_2_O. Amplicons were pooled at equimolar ratio. The pooled products were end-repaired by addition of 2.8 µl NEB Ultra II End Repair buffer and 1.2 µl NEB Ultra II End Repair Enzyme Mix, followed by incubation at RT for 5 min and 65 °C for 5 min. The end-repaired DNA was then purified and concentrated to 5 µl by SPRI beads (Omega Biotek MagBind NGS), followed by addition of 1.5 µl of native adapter ligation barcode (SQK-NBD114.96, ONT) and 6.5 µl of Blunt/TA ligation master mix (NEB). After a 20 min incubation at RT 1 µl EDTA (SQK-NBD114.96) was added, barcoded amplicons were pooled with other, unrelated samples containing different barcodes, the DNA was purified with 0.4 volumes of SPRI beads, washed once with short fragment buffer (SFB, SQK-NBD114.96) and twice with 80% EtOH before elution in 30 µl H_2_O. The pooled library was then subjected to motor protein ligation by addition of 10 µl NEB 5x Quick Ligation Buffer, 5 µl Kit 14 native adapter (NA, SQK-NBD114.96) and 5 µl of high concentration T4 DNA Ligase (NEB). The ligation reaction was incubated for 20 min at RT before purification with 0.4 volumes of Ampure XP beads (SQK-NBD114.96), washed and eluted in 15 µl elution buffer (EB, SQK-NBD114.96) at 37 °C for 10 min. The resulting library was quantified with the AccuClear Ultra High Sensitivity dsDNA Quantitation Solution (Biotium, Cat. No. 31027) and 10 fmol were loaded onto a Flongle R10.4.1 flow cell (ONT, FLO-FLG114) until the flow cell reached its end of life. Data was basecalled with model r10.4.1_e8.2_400bps_sup 5 kHz) with guppy version 6.5.7. Basecalled reads were then aligned to reference sequence NC_006579 of PVM strain J3666 using minimap2 and alignments were visualized with Intergrative Genomics Viewer (IGV version 2.19.4). WhatsHap (version 2.2) based phasing analysis to separate the two populations was performed using the FA-NIVA pipeline (Neurgaonkar et al., 2025). Per position mismatch rates were extracted with perbase version 0.8.5.

### Generation of rPVM mutants encoding the G gene variants of PVM J3666

The generation of recombinant PVM strain 15 (rPVM) has been described previously (18). Exchanges of the determining parts of the G gene were performed using a plasmid containing the antigenomic cDNA fragment encoding the complete G, F and M2 genes flanked by *Age*I and *BstB*I sites. This fragment corresponds to the fragment 3 used to assemble the original full-length cDNA clone (18). Fragments of corresponding J3666-G clones containing the 65U or 65A nucleotide, respectively, as well as other distinctive polymorphic nucleotides, were amplified using a forward primer (5-<u>ACCGGT</u>AGGATAATCTACCTATTG; *AgeI* site is underlined)) binding to the G gene start signal that would also add an *Age*I restriction site upstream of G that is unique for rPVM, and a reverse primer (5- TCCACTGCACTACTATAGATTGC) that binds 161 nucleotides downstream of the G gene in F. A fragment of the strain 15-G encompassing nt 1-931 was replaced with the equivalent portion from the G_J3666_65U amplificate using the *AgeI* and a natural occuring *AvrII* site. For replacement of the strain 15-G with G_J3666_65A, the entire G gene and the next 155 nucleotides downstream of the G gene were exchanged via *AgeI* and natural occurring *NdeI* sites (Fig. 3A). The resulting GJ3666-F-M2 fragments were used to exchange the corresponding portion of the original full-length cDNA clone pPVM via *Age*I and *BstB*I to obtain pPVM-G_J3666_65U and pPVM-G_J3666_65A respectively. Recombinant PVM-G_J3666_65U and rPVM-G_J3666_65A were then recovered as previously described (18). The sequences of replaced portions were confirmed before virus recovery and in the final virus stock after recovery as described above.

### Western blot analysis

Total protein lysates were prepared from BHK-21 cells infected at MOI of 0.1 per cell for 4 days. Cells were washed twice, collected with cell scrapper into ice-cold phosphate buffered saline (PBS). The cells were dissolved into 150 µl of PBS and lysed with equal amount of 2X Laemmli buffer (39). The lysates were homogenized using QIAshredder (Qiagen) and the total protein yields were quantified using the Roti-Quant universal kit (Roth, Karlsruhe Germany). Equal amounts of protein were separated on 8% SDS-containing polyacrylamide gels and transferred onto Protan blotting membrane (Whatmann GE, Freiburg Germany). Following blocking with 5% non-fat milk powder in PBS PVM-specific proteins were detected with polyclonal serum collected from a convalescent rPVM-infected mouse combined with a secondary goat anti-mouse HRP-conjugated antibody (Millipore, Darmstadt Germany). Images were recorded with the LAS 3000 CCD camera (Fujifilm). Two different blots from two similar infections were quantified using the AIDA software (Raytest Straubenhardt, Germany) and the results were analyzed with GraphPad prism 4.0. (GraphPad, La Jolla, USA).

### Animal experiments

Six to nine weeks old BALB/c mice were intranasally infected with a sublethal dose (150 PFU in 80 µl of phosphate-buffered saline) of virus suspension under isoflourane anesthesia. Daily weight estimation was used to monitor virulence of virus. For determination of virus titers, mice were sacrificed at day 3 and 6 post infection, the lungs removed and homogenized in 3 ml of EMEM containing 10% FCS, 50 mM HEPES and 100 mM MgSO_4_. The clarified homogenates were used for virus titration on Vero cells. All animal experiments were performed in conformity with the guidelines of the animal ethical committee of the University of Wuerzburg and the city of Wuerzburg, Germany (Az. 55.2-2532-2-318).

### Statistical analysis

Statistical analyses were performed with GraphPad prism 4.0 using appropriate test and an alpha-value of 0.05 to establish statistical significance, unless otherwise stated.

## Acknowledgements

Financial support was provided by the University of Würzburg Graduate School of Life Science (GSLS) to AA, and the Helmholtz Association VH-NG-1347 to RPS. RPS also acknowledges the interdisciplinary Thematic Institute IMCBio+, as part of the ITI 2021–2028 program of the University of Strasbourg, CNRS and Inserm, IdEx Unistra (ANR-10-IDEX-0002), SFRI-STRAT′US (ANR 20-SFRI-0012), and EUR IMCBio (ANR-17-EURE-0023) under the framework of the French Investments of the France 2030 Program. We thank Theresa Kreuzahler for excellent technical assistance.

## Author contributions

CDK and ARA conceptualized the project. ARA, MF and PB carried out experiments. PB, JY, AKB and RPS curated and analyzed data sets. AKB, RPS and CDK supervised experiments. ARA, PB, JY and CDK wrote the original manuscript. All authors reviewed and edited the manuscript.

---

**Supplementary figure S1.**
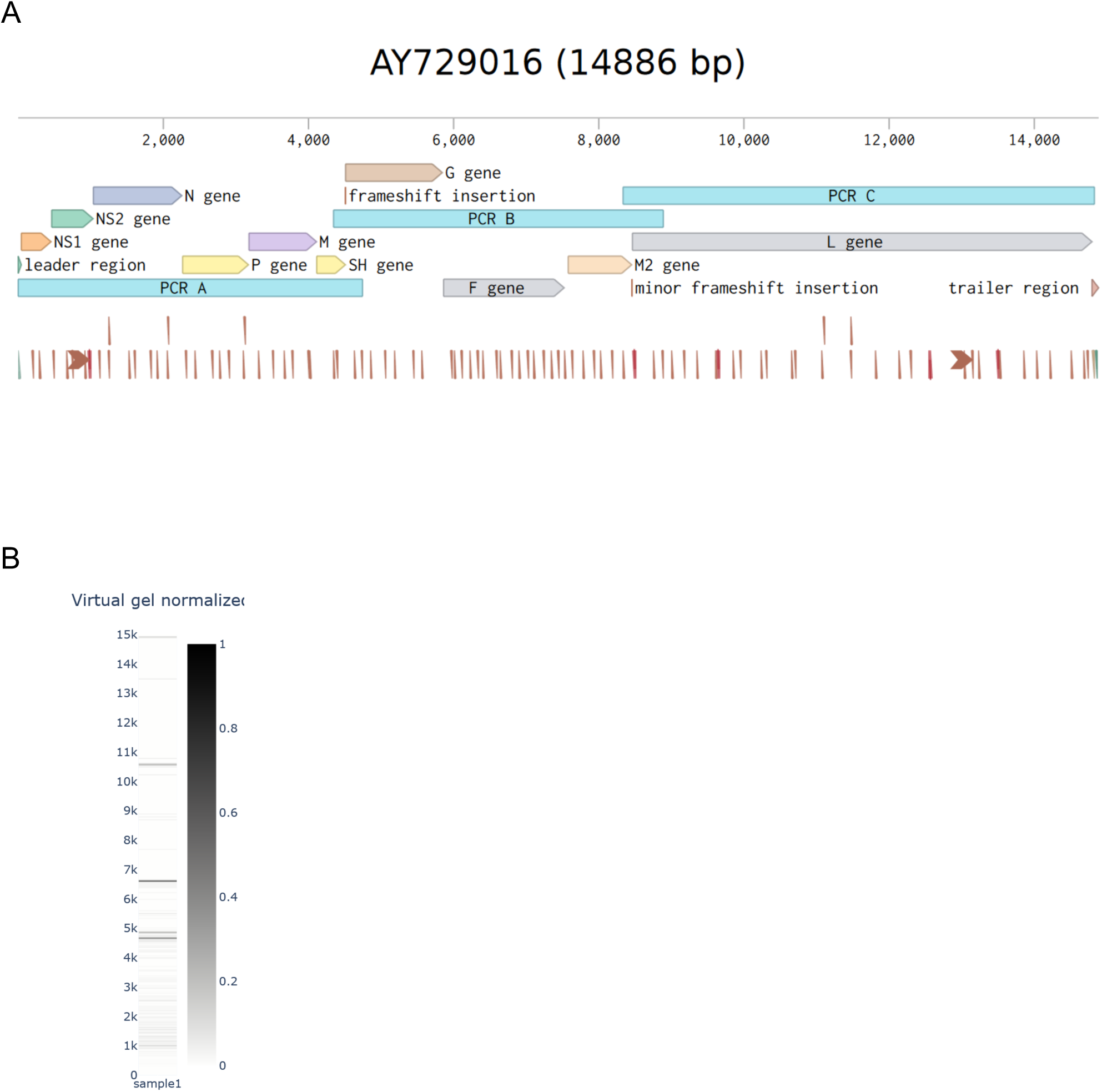
Oxford nanopore sequencing strategy. (A) PCR strategy for amplification of the PVM genome. The strategy was originally designed on the basis of PVM strain 15 that, however, has no relevance for the success. (B) Virtual gel of sequenced reads. Shown is the density of reads (with intensity scaled by number basepairs) across the 0 to 15000 nt read lengths.

**Supplementary figure S2.**
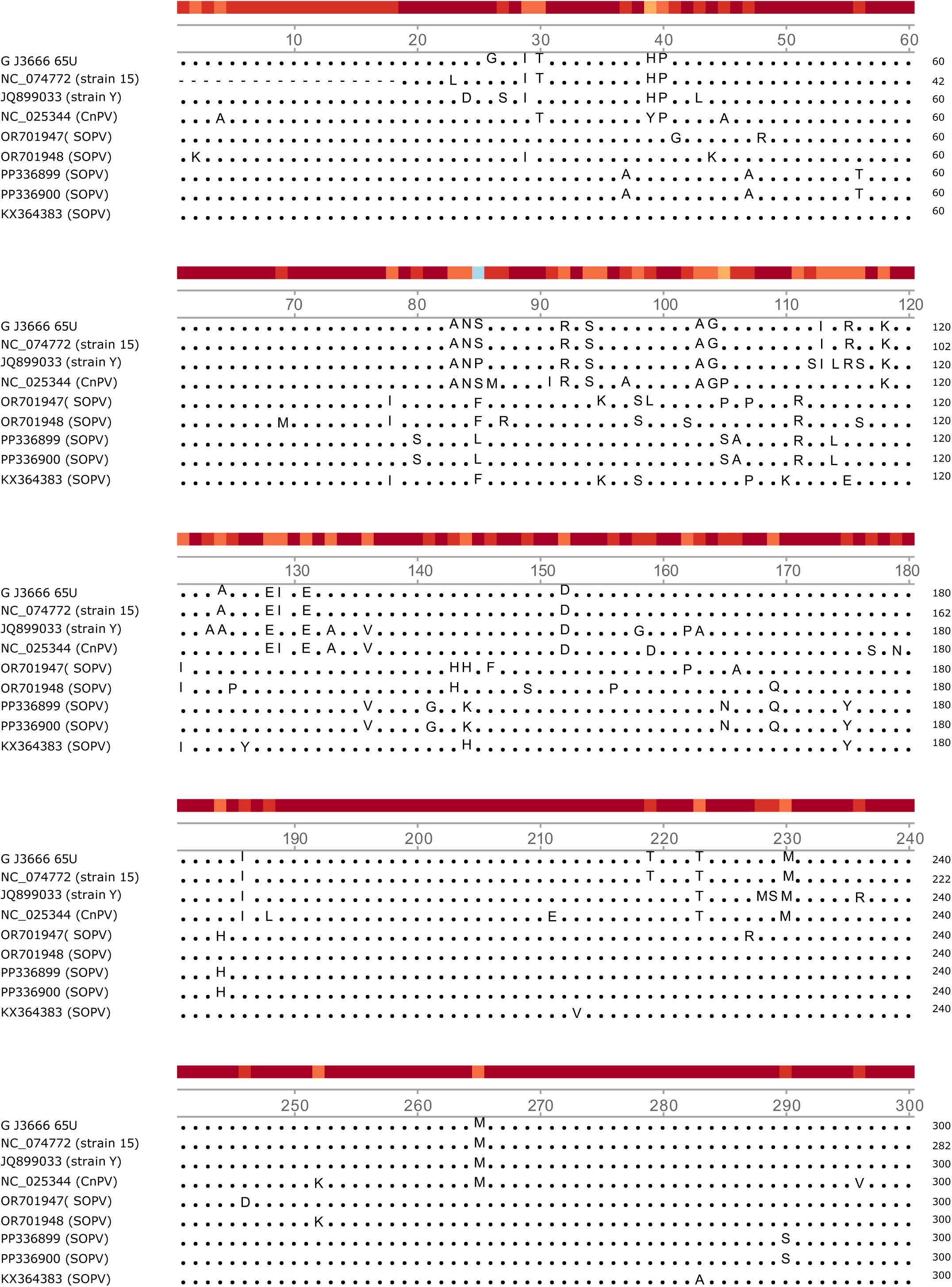
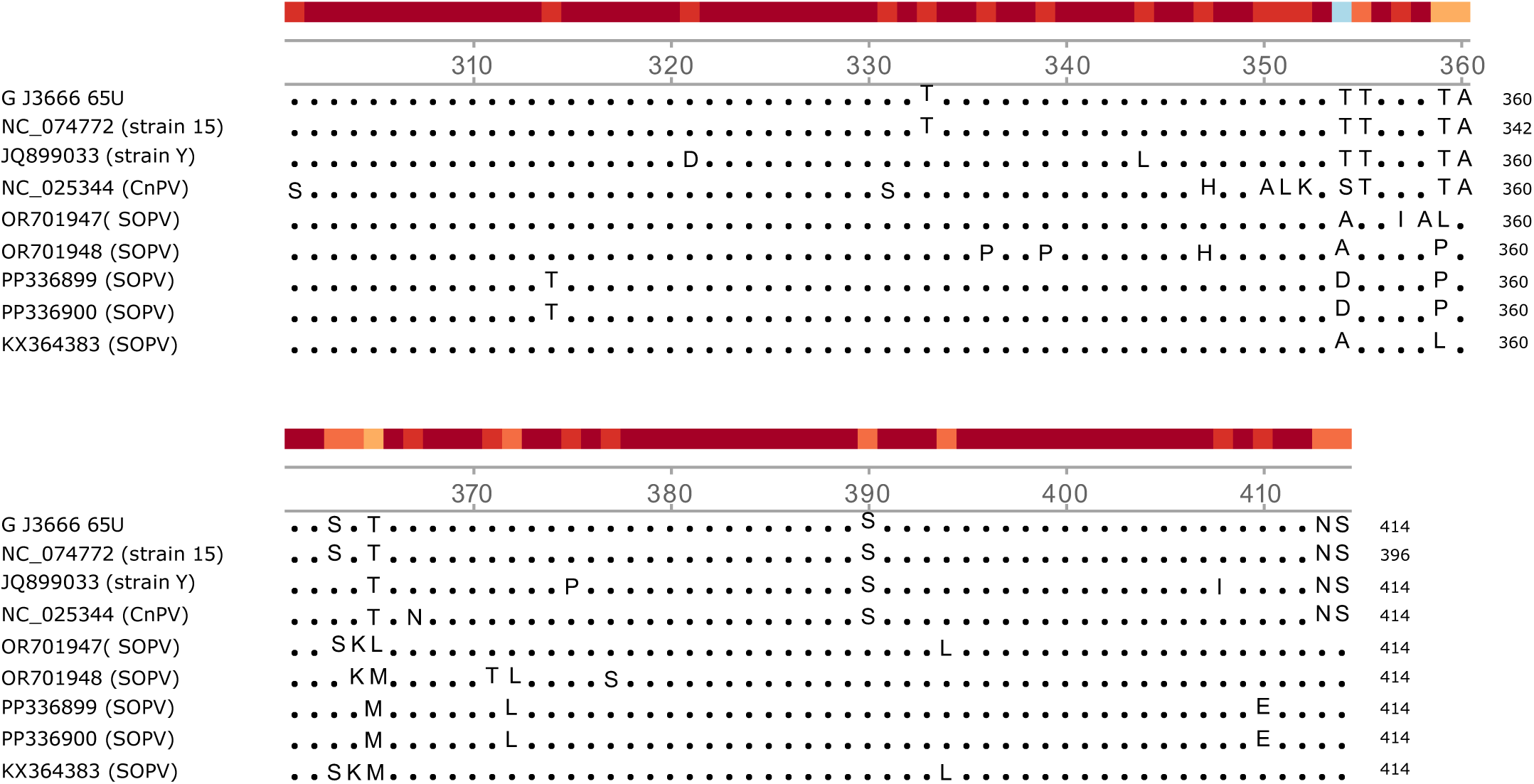
Alignment of PVM-related pneumovirus G proteins. Except for the first sequence which represents that of G J3666 with 414 amino acids described here, all sequences are identified by the GeneBank accession number and the species. The first three sequences belong to PVM G proteins.

**Supplementary figure S3.**
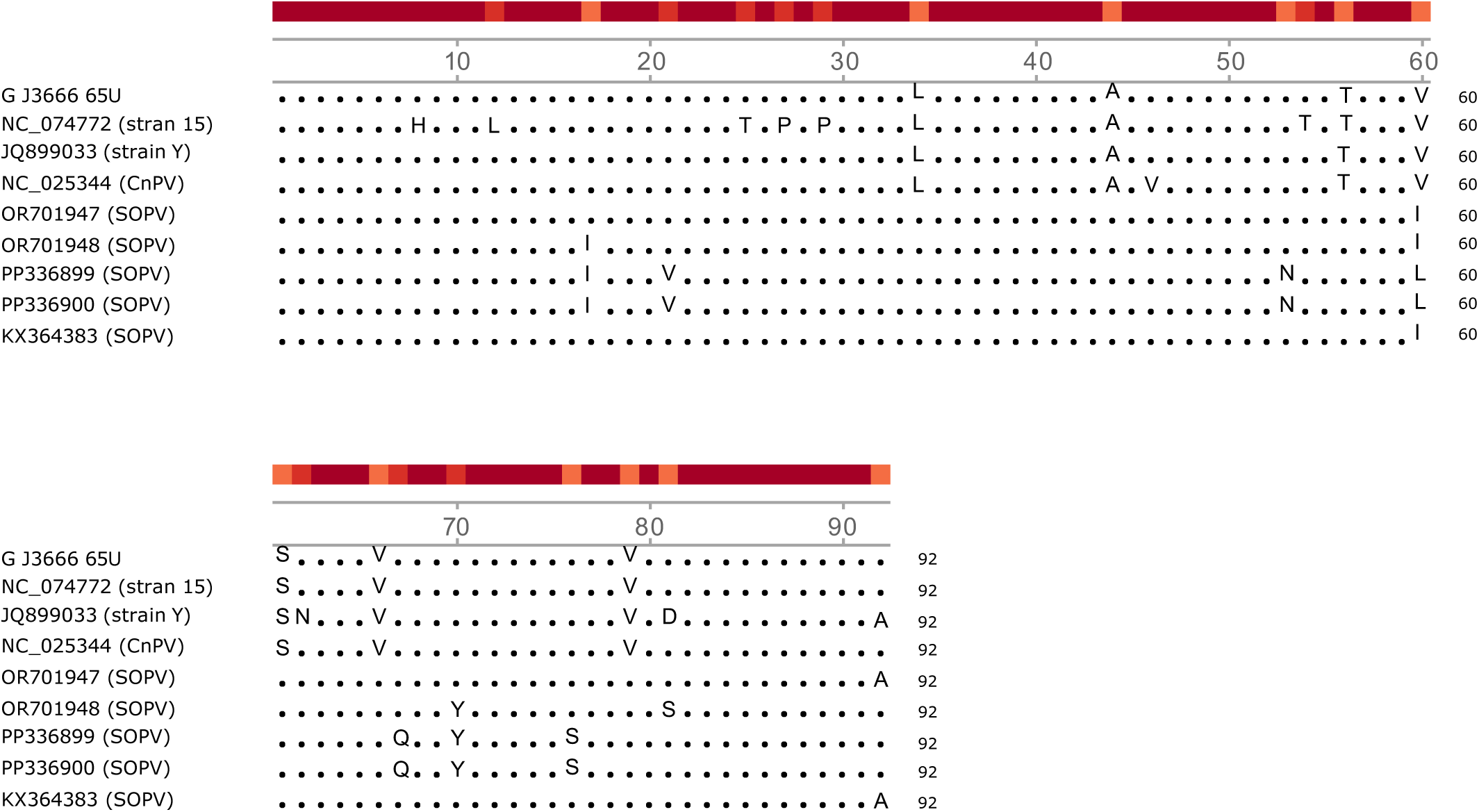
Alignment of PVM-related pneumovirus SH proteins. Except for the first sequence which represents that of SH J3666 with 92 amino acids described here, all sequences are identified by the GeneBank accession number and the species. The first three sequences belong to PVM SH proteins.

## References

1. Rima B, Collins P, Easton A, Fouchier R, Kurath G, Lamb RA, Lee B, Maisner A, Rota P, Wang L, Ictv Report C. 2017. ICTV Virus Taxonomy Profile: Pneumoviridae. J Gen Virol 98:2912–2913.

2. Horsfall FL, Hahn RG. 1940. A Latent Virus in Normal Mice Capable of Producing Pneumonia in Its Natural Host. J Exp Med 71:391–408.

3. Dyer KD, Garcia-Crespo KE, Glineur S, Domachowske JB, Rosenberg HF. 2012. The Pneumonia Virus of Mice (PVM) model of acute respiratory infection. Viruses 4:3494–510.

4. Weir EC, Brownstein DG, Smith AL, Johnson EA. 1988. Respiratory disease and wasting in athymic mice infected with pneumonia virus of mice. Lab Anim Sci 38:133–7.

5. Drexler JF, Corman VM, Muller MA, Maganga GD, Vallo P, Binger T, Gloza-Rausch F, Cottontail VM, Rasche A, Yordanov S, Seebens A, Knornschild M, Oppong S, Adu Sarkodie Y, Pongombo C, Lukashev AN, Schmidt-Chanasit J, Stocker A, Carneiro AJ, Erbar S, Maisner A, Fronhoffs F, Buettner R, Kalko EK, Kruppa T, Franke CR, Kallies R, Yandoko ER, Herrler G, Reusken C, Hassanin A, Kruger DH, Matthee S, Ulrich RG, Leroy EM, Drosten C. 2012. Bats host major mammalian paramyxoviruses. Nat Commun 3:796.

6. He WT, Hou X, Zhao J, Sun J, He H, Si W, Wang J, Jiang Z, Yan Z, Xing G, Lu M, Suchard MA, Ji X, Gong W, He B, Li J, Lemey P, Guo D, Tu C, Holmes EC, Shi M, Su S. 2022. Virome characterization of game animals in China reveals a spectrum of emerging pathogens. Cell 185:1117–1129 e8.

7. Glineur SF, Renshaw RW, Percopo CM, Dyer KD, Dubovi EJ, Domachowske JB, Rosenberg HF. 2013. Novel pneumoviruses (PnVs): Evolution and inflammatory pathology. Virology 443:257–64.

8. Hause BM, Padmanabhan A, Pedersen K, Gidlewski T. 2016. Feral swine virome is dominated by single-stranded DNA viruses and contains a novel Orthopneumovirus which circulates both in feral and domestic swine. J Gen Virol 97:2090–2095.

9. Park J, Kim HR, Lee EB, Lee SK, Kim WI, Lyoo YS, Park CK, Ku BK, Jeoung HY, Lee KK, Park SC. 2023. First Detection and Genetic Characterization of Swine Orthopneumovirus from Domestic Pig Farms in the Republic of Korea. Viruses 15.

10. Randhawa JS, Chambers P, Pringle CR, Easton AJ. 1995. Nucleotide sequences of the genes encoding the putative attachment glycoprotein (G) of mouse and tissue culture-passaged strains of pneumonia virus of mice. Virology 207:240–5.

11. Krempl CD, Collins PL. 2004. Reevaluation of the virulence of prototypic strain 15 of pneumonia virus of mice. J Virol 78:13362–5.

12. Thorpe LC, Easton AJ. 2005. Genome sequence of the non-pathogenic strain 15 of pneumonia virus of mice and comparison with the genome of the pathogenic strain J3666. J Gen Virol 86:159–169.

13. Cook PM, Eglin RP, Easton AJ. 1998. Pathogenesis of pneumovirus infections in mice: detection of pneumonia virus of mice and human respiratory syncytial virus mRNA in lungs of infected mice by in situ hybridization. J Gen Virol 79 ( Pt 10):2411–7.

14. Krempl CD, Lamirande EW, Collins PL. 2005. Complete sequence of the RNA genome of pneumonia virus of mice (PVM). Virus Genes 30:237–49.

15. Buchholz UJ, Ward JM, Lamirande EW, Heinze B, Krempl CD, Collins PL. 2009. Deletion of nonstructural proteins NS1 and NS2 from pneumonia virus of mice attenuates viral replication and reduces pulmonary cytokine expression and disease. J Virol 83:1969–80.

16. Heinze B, Frey S, Mordstein M, Schmitt-Graff A, Ehl S, Buchholz UJ, Collins PL, Staeheli P, Krempl CD. 2011. Both nonstructural proteins NS1 and NS2 of pneumonia virus of mice are inhibitors of the interferon type I and type III responses in vivo. J Virol 85:4071–84.

17. Dibben O, Thorpe LC, Easton AJ. 2008. Roles of the PVM M2-1, M2-2 and P gene ORF 2 (P-2) proteins in viral replication. Virus Res 131:47–53.

18. Krempl CD, Wnekowicz A, Lamirande EW, Nayebagha G, Collins PL, Buchholz UJ. 2007. Identification of a novel virulence factor in recombinant pneumonia virus of mice. J Virol 81:9490–501.

19. Buchholz UJ, Anderson LJ, Collins PL, Mejias A. 2023. Respiratory Syncytial Virus and Metapneumovirus, p 267 - 317. *In* Howley PM, Knipe DM (ed), Fields Virology, 7 ed, vol 3. Wolters Kluwer, Philadelphia, PA.

20. Karron RA, Buonagurio DA, Georgiu AF, Whitehead SS, Adamus JE, Clements-Mann ML, Harris DO, Randolph VB, Udem SA, Murphy BR, Sidhu MS. 1997. Respiratory syncytial virus (RSV) SH and G proteins are not essential for viral replication in vitro: clinical evaluation and molecular characterization of a cold-passaged, attenuated RSV subgroup B mutant. Proc Natl Acad Sci U S A 94:13961–6.

21. Techaarpornkul S, Barretto N, Peeples ME. 2001. Functional analysis of recombinant respiratory syncytial virus deletion mutants lacking the small hydrophobic and/or attachment glycoprotein gene. J Virol 75:6825–34.

22. Teng MN, Whitehead SS, Collins PL. 2001. Contribution of the respiratory syncytial virus G glycoprotein and its secreted and membrane-bound forms to virus replication in vitro and in vivo. Virology 289:283–96.

23. Renshaw RW, Zylich NC, Laverack MA, Glaser AL, Dubovi EJ. 2010. Pneumovirus in dogs with acute respiratory disease. Emerg Infect Dis 16:993–5.

24. Renshaw R, Laverack M, Zylich N, Glaser A, Dubovi E. 2011. Genomic analysis of a pneumovirus isolated from dogs with acute respiratory disease. Vet Microbiol 150:88–95.

25. Decaro N, Pinto P, Mari V, Elia G, Larocca V, Camero M, Terio V, Losurdo M, Martella V, Buonavoglia C. 2014. Full-genome analysis of a canine pneumovirus causing acute respiratory disease in dogs, Italy. PLoS One 9:e85220.

26. Piewbang C, Techangamsuwan S. 2019. Phylogenetic evidence of a novel lineage of canine pneumovirus and a naturally recombinant strain isolated from dogs with respiratory illness in Thailand. BMC Vet Res 15:300.

27. Song X, Li Y, Huang J, Cao H, Zhou Q, Sha X, Zhang B. 2021. An emerging orthopneumovirus detected from dogs with canine infectious respiratory disease in China. Transbound Emerg Dis 68:3217–3221.

28. Dunowska M, More GD, Biggs PJ, Cave NJ. 2024. Genomic analysis of canine pneumoviruses and canine respiratory coronavirus from New Zealand. N Z Vet J 72:191–200.

29. Thieulent CJ, Carossino M, Peak L, Wolfson W, Li G, Balasuriya UBR. 2024. Coding-complete genome sequences of two strains of canine pneumovirus derived from dogs with upper respiratory disease in the United States. Microbiol Resour Announc 13:e0105723.

30. Richard CA, Hervet C, Ménard D, Gutsche I, Normand V, Renois F, Meurens F, Eléouët JF. 2018. First demonstration of the circulation of a pneumovirus in French pigs by detection of anti-swine orthopneumovirus nucleoprotein antibodies. Vet Res 49:118.

31. Graaf-Rau A, Hennig C, Lillie-Jaschniski K, Koechling M, Stadler J, Boehmer J, Ripp U, Pohlmann A, Schwarz BA, Beer M, Harder T. 2023. Emergence of swine influenza A virus, porcine respirovirus 1 and swine orthopneumovirus in porcine respiratory disease in Germany. Emerg Microbes Infect 12:2239938.

32. Ellis JA, Martin BV, Waldner C, Dyer KD, Domachowske JB, Rosenberg HF. 2007. Mucosal inoculation with an attenuated mouse pneumovirus strain protects against virulent challenge in wild type and interferon-gamma receptor deficient mice. Vaccine 25:1085–95.

33. Dever TE, Ivanov IP, Hinnebusch AG. 2023. Translational regulation by uORFs and start codon selection stringency. Genes Dev 37:474–489.

34. Kozak M. 1984. Point mutations close to the AUG initiator codon affect the efficiency of translation of rat preproinsulin in vivo. Nature 308:241–246.

35. Buchholz UJ, Finke S, Conzelmann KK. 1999. Generation of bovine respiratory syncytial virus (BRSV) from cDNA: BRSV NS2 is not essential for virus replication in tissue culture, and the human RSV leader region acts as a functional BRSV genome promoter. J Virol 73:251–9.

36. Marino JH, Cook P, Miller KS. 2003. Accurate and statistically verified quantification of relative mRNA abundances using SYBR Green I and real-time RT-PCR. J Immunol Methods 283:291–306.

37. Ruijter JM, Ramakers C, Hoogaars WM, Karlen Y, Bakker O, van den Hoff MJ, Moorman AF. 2009. Amplification efficiency: linking baseline and bias in the analysis of quantitative PCR data. Nucleic Acids Res 37:e45.

38. Zhao C, Liu F, Pyle AM. 2018. An ultraprocessive, accurate reverse transcriptase encoded by a metazoan group II intron. RNA 24:183–195.

39. Laemmli UK. 1970. Cleavage of structural proteins during the assembly of the head of bacteriophage T4. Nature 227:680–5.

